# F.O.R.W.A.R.D: A Data-Driven Framework for Network-Based Target Prioritization in Drug Discovery

**DOI:** 10.1101/2024.07.16.602603

**Authors:** Saptarshi Sinha, Ella McLaren, Madhubanti Mullick, Harrison M. Penrose, Courtney Tindle, Siddharth Singh, Brigid S. Boland, Pradipta Ghosh

## Abstract

Despite advances in artificial intelligence (AI), target-based drug development remains costly, complex, and imprecise. We introduce F.O.R.W.A.R.D [*Framework for Outcome-based Research and Drug Development*], a network-based target prioritization platform, and demonstrate its utility in the challenging landscape of Inflammatory Bowel Diseases (IBD), a chronic, multifactorial condition. F.O.R.W.A.R.D uses real-world clinical outcomes, and a machine-learning classifier trained on transcriptomic data from seven prospective randomized trials across four drugs. It defines remission at the molecular level and calculates, using network connectivity, the likelihood that targeting a given molecule will induce remission-associated gene expression. Benchmarking against 210 completed trials across 52 targets, F.O.R.W.A.R.D achieved 100% predictive accuracy—despite variability in drug mechanisms and trial designs. Single-cell RNA-seq and a prospective biobank of patient-derived organoids confirmed that the remission signature is epithelium-specific and tracks with poor outcomes. F.O.R.W.A.R.D enables *in-silico* phase zero trials to inform trial design, revive shelved drugs, and guide early termination decisions. Broadly applicable and iteratively refined by emerging trial data, F.O.R.W.A.R.D has the potential to reshape drug discovery—bringing foresight to hindsight, and empowering both R&D and human-in-the-loop clinical decision-making.

## Introduction

The average drug takes 10–12 years and $1–1.8 billion to progress through clinical trials to Food and Drug Administration (FDA) approval^1^. The efficiency of drug development has steadily declined since the 1950s; with a ∼90% failure rate^2^, the cost of bringing a new drug to market doubles roughly every nine years—a trend dubbed “Eroom’s Law”^3,4^. This is a playful inversion of Moore’s Law (a measure of increasing computing efficiency^5^), highlighting a paradox. This paradox persists despite the rise of precision medicine^6,7^ and rapid advances in artificial intelligence (AI), which promised to accelerate discovery by harnessing big data.

In addition, AI’s impact on drug development remains in question^6,8,9^, especially in three key areas ^8^: (i) *Target discovery —many “AI-discovered” targets were already known, raising doubts about novelty;* (ii) *Success rates* —AI-driven trials show similar phase 1/2 success rates (∼87% and ∼40%, respectively ^10^) to non-AI programs (∼75% and ∼50% ^11^), with phase 3 results still pending; and (iii) *Process innovation —*most AI tools (especially large language models; LLMs) rely on conventional approaches like literature mining and omics integration, offering limited advancement in methodology.

Thus, a critical unmet need remains: improving precision in target selection, which underlies the two main reasons for failure—lack of efficacy (50–60%) and safety/toxicity issues (22–35%)^12–14^.

Here, we present a machine learning–based framework that integrates diverse transcriptomic datasets linked to clinical outcomes to define a remission-associated gene signature—one that appears uniquely specific to the gut epithelium. We compute network-based connectivity between this signature and candidate drug targets to guide prioritization and validate key predictions using prospective studies with patient-derived organoids (PDOs). This approach yields a robust, accurate model that identifies core cellular and molecular determinants of therapeutic response and can be integrated into existing pipelines as a modular decision-support tool. By enhancing precision in early-stage target selection, this framework offers the potential to significantly improve R&D efficiency and reshape the landscape of drug discovery.

## Materials and Methods

### Boolean representation of gene expression data

Boolean logic represents the mathematical relationship between two values, such as high(1)/low(0). To perform Boolean analysis on gene expression data, the expression levels of the two comparator genes must first be converted into one of the two possible values (0/1). Such conversion is done using the StepMiner^15^ algorithm which determines a threshold considering the entire dynamic range of the gene’s expression profile across the entire sample set. This algorithm employs adaptive regression to identify the optimal fit for transitions, whether from high to low or low to high. A noise margin corresponding to a 2-fold change is applied around the threshold to identify intermediate values, which are disregarded during Boolean analysis. The relationship between two genes, i.e., drug’s target gene (T) and one of the 34 reference genes within the molecular signature of disease remission (R), can be plotted as a scatter plot in each set of samples. In such scatterplots, the x and y-axis represent the StepMiner normalized expression values of R and T genes, respectively (**Fig S1A**). Based on the StepMiner thresholds of both genes, four possible quadrants can be identified in any scatter plot. The statistical sparsity (as determined by *Boolean Implication statistics*; see next) of sample distribution in any of these quadrants enables the estimation of Boolean Implication Relationships (BIRs). These calculations were carried out in the Boolean Network Explorer (BoNE^16^), a computational tool designed to analyze and visualize Boolean networks which carries the statistical modules for the estimation of binary states, BIRs, and carrying out analyses using adaptive regression.

### Boolean implication statistics

Boolean implication relationship between reference gene, R, and drug’s target gene, T, can be computed based on the sample distribution in the four quadrants (0, 1, 2 and 3) based on the normalized values of R and T genes in each sample (**Fig S1A**). A Boolean implication relationship is identified if any one of the four possible quadrants or two diagonally opposite quadrants are sparsely populated. Based on this criterion, there are six different types of Boolean implication relationships^15^ (**Fig S1B**). Two of these are symmetric: equivalent (corresponding to highly positively correlated genes, if quadrants 1 and 2 are sparse) and opposite (corresponding to highly negatively correlated genes, if quadrants 0 and 3 are sparse). The other four Boolean relationships are asymmetric, each corresponding to one sparsely populated quadrant: (T low => R low, if quadrant 2 is sparse), (T high => R low, if quadrant 3 is sparse), (T low => R high, if quadrant 0 is sparse), and (T high => R high, if quadrant 1 is sparse).

The sparse quadrant can be identified using BooleanNet statistics (*S*) and error rate (*p*). Considering the total number of samples under is *N* (*N* = *n*_0_ + *n*_1_ + *n*_2_ + *n*_3_), where *n*_0_, *n*_1_, *n*_2_ and *n*_3_ are the number of samples in 0, 1, 2 and 3 quadrants respectively (**Fig S1A**). Based on StepMiner threshold, the number of samples with low R values will be (*n*_0_ + *n*_1_), whereas the number of samples with low T values will be (*n*_0_ + *n*_2_). So, the observed number of samples in quadrant 0 is *n*_0_, the expected number of samples in quadrant 0 (*n*^′^) will be computed as:

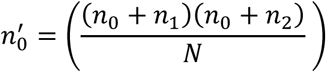

So, the sparsity of quadrant 0 can be calculated as,

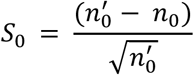

And the error rate can be estimated as,

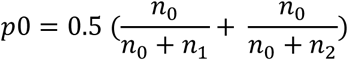

### Computing the therapeutic index (TI) of target gene(s)

The therapeutic index (TI) is a computationally generated metric based on Boolean implication statistics that provides a likelihood score indicating how closely the proposed target gene (T) is associated with a given reference gene (R). A quadrant is considered as sparse if *S* is high (*n*^′^ > *n*) and p is small. A quadrant is considered sparse when *S* > 1 and *p* < 0.1^17^. There is a key assumption in this approach that ties a drug’s mechanism of action to the gene’s expression: drugs that target gene(s) as *ago*nists, are presumed to upregulate the expression of gene T. Similarly, drugs that target gene(s) as *anta*gonists, are presumed to suppress the expression of gene T. This set of assumptions is no different than what is currently used in conventional drug discovery approaches which consider inhibitors for upregulated candidate genes in a set of DEGs, and activators for downregulated candidate genes in a set of DEGs.

The TI of the target genes that are *ago*nistically targeted will be computed based on their co-expression relationships with the reference gene, i.e., ‘T equivalent R’ and ‘T high=> R high’ (**Fig S1C-D**). Simply put, this means that when the expression of gene T is induced by the drug, the probabilities are high that the expression of R is co-induced through the fundamental gen regulatory processes that are captured by the BIRs. To increase the rigor, we ignore the ‘T high => R low’ relationship; this is because in this instance, if T is induced, there is a probability that R might not be co-induced. If T is *ago*nistically targeted, the TI of T will be computed as:

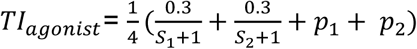

Similarly, the TI of the target genes that are *anta*gonistically targeted will be computed based on their opposing expression relationships with the reference gene, i.e., ‘T opposite R’ and ‘T low=> R high’ (**Fig S1E**). Simply put, this means that when the expression of gene T is suppressed by the drug, the probabilities are high that R is induced through the fundamental gene regulatory processes that are captured by the BIRs. To increase the rigor, we ignore the ‘T high => R low’ relationship; this is because in this instance, if T is suppressed, there is a probability that R might not be induced. If T is *anta*gonistically targeted, the TI of T will be computed as:

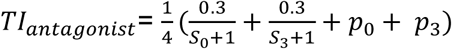

In other words, if the drug target is *ago*nistically targeted then the Boolean implication relation of the drug target with the reference gene should have the *S*_1_ > 1. Conversely, if the drug target is *anta*gonistically targeted then the Boolean implication relation of the drug target with the reference gene should have the *S*_0_ > 1 (**Fig S1F**). Lower the TI, the stronger the association between T and R. Here, we have considered the association between T and R if the TI is <0.075, optimized based on the target genes of the four FDA-approved drugs.

### Connectivity index (CI) and Likelihood of approval index (LoAI)

The connectivity index (CI) of a drug target is a fractional value that indicates how many of the 34 reference genes (R) are strongly associated with the drug target gene (T), as determined by Boolean implication statistics. Specifically, the CI for a given target gene (T gene) is calculated as the fraction of reference genes (R genes) that have a TI (Boolean implication statistic) value less than 0.075 with the T gene.

The Likelihood of Approval Index (LoAI) measures the overall impact of a drug by combining the effects of all the genes whose expression it alters. For example, for drugs that target protein(s) encoded by single genes, the CI = LoAI. However, for drugs that target proteins encoded by multiple genes, the LoAI is a union (ᵁ) of the CI of the individual genes. That way, LoAI ensures that when a drug has multiple target genes, each target’s connectivity to the reference genes is counted only once, thereby avoiding redundancy. In doing so, LoAI results in a comprehensive score that reflects the drug’s potential effectiveness to predictably induce the majority of the 34 reference genes of remission. In drugs with multiple target genes, there were two other possibilities considered: (i) some target genes of the same drug are predicted to either induce the 34 reference genes (intended use CI), while others are not connected at all to the reference genes (i.e., >/= 0.075 TI). (ii) some of the target genes are predicted to induce the 34 reference genes (intended use CI), whereas other target genes are connected in ways that are predicted to suppress the 34 reference genes (unintended use CI). In the first instance, the LoAI is a comprehensive score of all intended use CI. In the second instance, the LoAI is computed by subtracting unintended from intended use CI.

### Cumulative connectivity (*C*_*T*_)

The cumulative connectivity (*C*_*T*_) of a drug quantifies the total number of reference genes associated with all its targets, including those that are redundant or overlapping. This measure could inform relative performance of two drugs that have crossed the threshold LoAI of >/=0.5 and may have some target genes shared between them. Given the reference gene set as G, the cumulative connectivity can be expressed as,

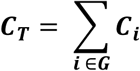

Where *C*_*i*_ indicates the connectivity of each reference gene, *i* (*i* ∈ *G*), with any of the target genes of the given drug.

### Composite score of expression of a set of genes

When calculating the composite score of the degree of expression of a set of genes, their expression levels in these datasets were first converted to binary values (high or low) using the StepMiner algorithm. Subsequently, their expression values were normalized (*Exp_St_*) using a modified Z-score approach, centered around the StepMiner threshold (*S_Thr_*) using the formula:

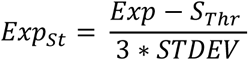

The normalized expression values for each gene were summed to create a final composite score for the gene signature. These composite scores reflect the overall activity or state of biological pathways associated with the genes, allowing for the identification of differences between control and query groups within any given dataset. However, composite scores cannot be directly compared between different gene signatures within a dataset, as they are not normalized by the number of genes in each signature. Additionally, the same signature cannot be compared across different datasets due to individual normalization based on intra-dataset sample distribution or other inherent differences. The samples were ordered based on their final signature score.

### Measurement of classification strength or prediction accuracy

To evaluate the classification strength and prediction accuracy of our gene expression–based models, we performed Receiver Operating Characteristic (ROC) analysis using the Python Scikit-learn package. ROC curves plot the True Positive Rate (TPR) against the False Positive Rate (FPR) at various threshold settings, and the Area Under the Curve (AUC) provides a measure of a classifier’s ability to discriminate between classes. AUC values closer to 1.0 indicate stronger discriminatory power, while values near 0.5 suggest performance no better than random chance. We selected a logistic regression–based classifier (rather than penalized regression methods) for initial gene-level evaluation. This is because penalized regression was found to underperform due to over-regularization, high feature collinearity, and class imbalance— limitations well-documented in high-dimensional biological data. For feature reduction, we employed univariate logistic regression for each of the 1284 genes, computing classification accuracy across all training datasets and selecting those with a mean AUC > 0.7. This threshold, though conservative, aligns with widely accepted conventions in biomarker discovery, where AUC ≥ 0.7 is considered indicative of moderate discriminatory ability^18^. The area under the ROC curve (AUC) measures the diagnostic ability of a binary classifier (e.g., high vs. low gene expression levels) as the discrimination threshold is varied. ROC curves plot the True Positive Rate (TPR) against the False Positive Rate (FPR) at different threshold settings. The AUC quantifies the likelihood that the classifier will correctly distinguish between randomly chosen samples from two groups.

While simpler than multivariate penalized approaches (e.g., LASSO, Ridge, or Elastic Net), univariate logistic regression combined with performance-based filtering remains a robust, interpretable, and widely adopted method in systems biology and transcriptomics^19^. This strategy avoids overfitting in high-dimensional data by evaluating each gene independently, allowing for biologically meaningful interpretation of individual gene–outcome relationships. Importantly, these univariate AUC values were not used directly for prediction; rather, they served to reduce the feature space before constructing the final multivariate predictive models.

To assess the parsimony and predictive strength of the reduced 34-gene model relative to the full 1284-gene set, we computed Likelihood Ratio test (LRT), Akaike Information Criterion (AIC) and Bayesian Information Criterion (BIC) using Python sklearn module.

### Analysis of single cell RNA seq datasets

Single-cell RNA-seq data from GSE214695 and GSE282122 were downloaded from the Gene Expression Omnibus (GEO) in H5AD format. The filtered barcode matrix data were processed using the Scanpy v1.10 package. Harmonypy was applied to perform batch normalization across samples. Cells were retained if they expressed a minimum of 5,000 genes, and genes were retained if they were expressed in at least 10 cells. Cell types were annotated using the ‘Cells_Intestinal_Tract’ model^20^ implemented in CellTypist v1.6.3. Pseudobulk datasets were generated by aggregating raw counts from individual cell subtypes, followed by normalization using the log2(CPM + 1) transformation. Downstream BoNE-based classification accuracy analyses were performed using the pseudobulk data in a manner consistent with standard bulk RNA-seq workflows.

### Validation on a prospective biobank of patient derived organoids (PDOs)

Previously established patient-derived organoids^21^ (PDOs) from patients with ulcerative colitis (UC) and Crohn’s disease (CD), molecularly phenotyped through multi-omics profiling^21^, were used to assess a 34-gene composite expression signature. Comprehensive clinical metadata were curated, including demographics (age, gender, ethnicity), disease classification (Montreal subtypes, B1-B3, history of strictures, prior ileocecal resection), disease activity indices (CD: Patient-Reported Outcomes [PRO], Simple Endoscopic Score for Crohn’s Disease [SES-CD]; UC: Partial Mayo Score [pMayo], Mayo Endoscopic Subscore [MES], Ulcerative Colitis Endoscopic Index of Severity [UCEIS]), biopsy location, therapeutic history (including use and response to advanced therapies), and PDO molecular subtype.

All procedures and subsequent access of medical records during follow up were conducted under IRB-approved protocols (UCSD Project ID# 190105, HUMANOID™ Center), in compliance with the Human Research Protection Program and HIPAA regulations. Participants provided written informed consent for colonic biopsy collection, PDO generation, and use of associated clinical data. Healthy controls were derived from histologically normal colonic tissue obtained during routine screening colonoscopies or evaluations for irritable bowel syndrome.

To enable cross-disease comparison, parallel clinical parameters (e.g., SES-CD vs. MES or UCEIS; PRO vs. pMayo) were normalized using min-max scaling. Univariate ordinary least squares (OLS) regression analyses were conducted via the Python *statsmodels* module to examine associations between the composite gene score and individual clinical variables. The p-values test the null hypothesis that each coefficient is equal to zero, indicating no effect.

## Results

### Study Design and Rationale

To integrate diverse datasets while minimizing the risk of overfitting, we employed a Boolean logic–based transcriptomic network approach grounded in two foundational concepts from mathematics and biology^22^. This framework, known as the Boolean Implication Network, models genes or gene clusters (nodes) connected by Boolean Implication Relationships (BIRs; directed edges). BIRs capture all six possible gene associations—two symmetric and four asymmetric^15^ —with the latter often overlooked by conventional methods. These invariant relationships are believed to represent fundamental gene regulatory events that persist across the continuum from health to disease^16,23–26^. Boolean Implication Networks have been used to generate predictive “disease maps,” which chart consistent, sequential gene expression changes found across all disease samples—regardless of clinical heterogeneity—reflecting continuum states between health and disease^15–17,24,26–30^. This robustness makes them particularly effective in capturing disease-defining signatures across diverse validation datasets. Comparative studies show that Boolean-derived signatures outperform conventional methods (e.g., differential expression, Bayesian inference, clustering), with lower risk of overfitting and greater relevance across patient populations ^15–17,24,26–30^. Unlike pathway enrichment–based models, this approach focuses directly on gene perturbations, avoiding biases that are introduced during functional annotation^31,32^.

While this framework is broadly applicable, we selected inflammatory bowel diseaseS (IBD) as a test case for three reasons. (i) IBD is a chronic, complex, heterogeneous autoimmune disorder shaped by non-intuitive interplay among the microbiome, genetics, environment, and immune system disease^33^, making it well suited for network-based modeling. (ii) IBD has a rich pipeline of past and ongoing drug development programs^34^, allowing retrospective benchmarking for predictive accuracy. De-risking phase 2 trials is an urgent unmet need in IBD^35^, as evidenced by costly late-stage failures like mongersen (anti-SMAD7) ^36,37^, and etrolizumab (β7-integrin) ^38^. (iii) A rigorously validated Boolean-network model of IBD is available^16^ (**Fig 1***-Step 1*); developed using ∼1500 unique IBD samples. This revealed universal downregulation of 1,284 genes, highlighting a consistent molecular signature of impaired gut barrier integrity and bioenergetics^16^.

**Figure 1.**
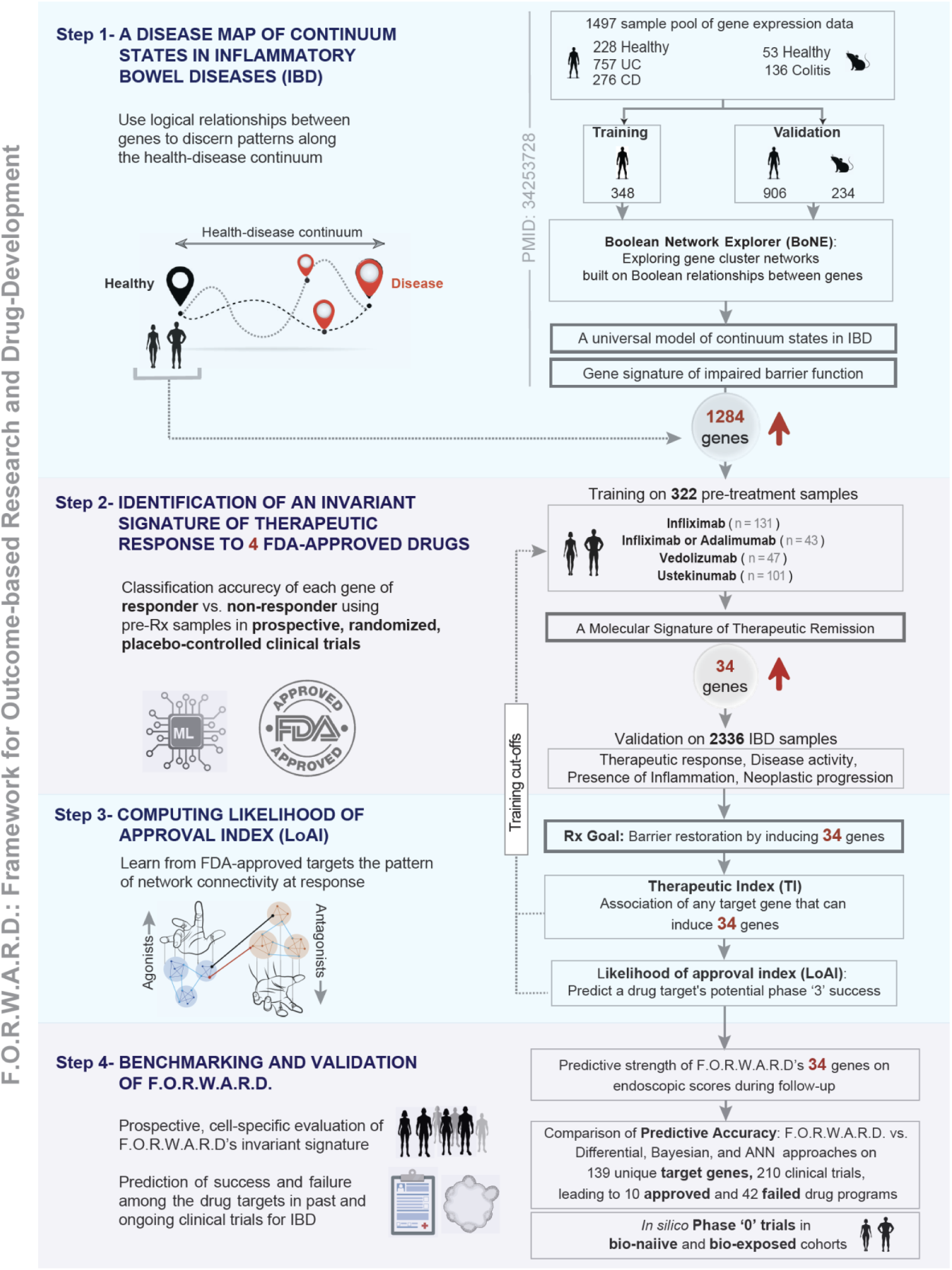
Study motivation, rationale and design. The ***four steps*** of a workflow that led to the development of a framework for outcome-based drug target prioritization (F.O.R.W.A.R.D) are presented. ***Step 1***: A published Boolean implication network-based computational model of disease continuum states in IBD ^16^, which serves as the premise of this study. The model was created and rigorously validated on transcriptomic databases containing overall 1497 gene–expression data (1263 human and 234 mouse samples). ***Step 2***: Use of a machine learning (ML) classifier to identify a 34-gene molecular signature of barrier restoration that is shared among all responders, regardless of therapeutic modality. Outcome-annotated datasets from real-world randomized, placebo-controlled prospective clinical trials that were used for training and validation are shown. A flowchart of the model training can be found in **Fig S2A. *Step 3***: Formulation of a Boolean implication-based connectivity index based on outcome-specific learning from real-world drug trials to standardize the model. Here, statistical learning is used to determine the logic and strength (TI) and degree (LoAI) of connectivity of a drug’s target gene (s) to the 34 genes. A LoAI >/= 0.50 was determined as likely to pass regulatory approval. ***Step 4***: Benchmarking and validation of F.O.R.W.A.R.D was done by three approaches: (i) prospective study to determine its cell-type specific abilities to predict endoscopic remission; (ii) test its predictive accuracy on drug development programs with known outcomes for the presence or lack of efficacy; (iii) test its ability to inform trial design through *in silico* trials on custom cohorts.

We trained a machine learning classifier on the published 1,284-gene full model using data from 7 clinical trial cohorts (n = 322 subjects; **Supplemental Datasheet 1**) to identify whether a shared therapeutic response signature exists, independent of drug class or target. Validation in 9 additional cohorts (n = 2336 subjects; **Supplemental Datasheet 1**) spanned various therapeutic modalities, disease states, and histopathological contexts (see **Fig 1***-Step 2*). A reduced 34-gene model emerged, whose reduction consistently marked lack of therapeutic response.

This inspired us to formalize the therapeutic goal as upregulation of these 34 genes through targeted perturbation of network-associated genes (**Fig 1***-Step 3*). Using principled network-based approaches, which included a combination of machine learning and statistical learning techniques, we computed the logic, strength, and connectivity of target-to-signature relationships within the Boolean network to inform rational drug discovery (**Fig 1***-Step 3*).

Finally, using single-cell RNA seq (scRNA Seq) and patient-derived organoids (PDOs), we pinpointed a surprisingly specific cellular and molecular feature that is the core determinant of therapeutic response and validated its ability to track endoscopic remission (**Fig 1***-Step 4*). We also benchmark the framework on 52 targets with known outcomes, compare its predictive accuracy against other approaches, and demonstrate its utility in *in-silico* phase 0 trials on custom cohorts (**Fig 1***-Step 4*).

### Identification of a molecular signature of clinical remission and therapeutic goal

We began with the full model of 1,284 genes, whose consistent downregulation across all IBD samples had previously been identified as indicative of epithelial barrier dysfunction^16^. Given that a compromised epithelial barrier is widely recognized as a key predictor of disease relapse, therapeutic response, and remission^39–44^, we asked whether specific aspects of this barrier defect might define remission. We trained the initial 1284-gene model by evaluating each gene’s ability to classify pre-treatment samples as responders or non-responders across seven prospective, randomized, placebo-controlled clinical trials involving four FDA-approved biologics (**Fig 2A**). Applying a stringent outcome-based training threshold (mean AUC 0.70 and standard deviation of 0.07) yielded a refined and reduced model of 34 genes (**Fig S2A**). A non-significant Likelihood Ratio Test (LRT) p-value (p > 0.05) indicates that the 34-gene model does not significantly underperform relative to the original 1,284-gene model (**Supplementary Table 1**). Additionally, lower Akaike Information Criterion (AIC) and Bayesian Information Criterion (BIC) values for the 34-gene model suggest it retains the essential predictive signal of the full model while minimizing overfitting (**Fig S2B-C**). The 34-gene signature consistently distinguished responders from non-responders across all training cohorts, regardless of the biologic therapy or its mechanism of action. In multivariate analyses of training cohorts with ≥3 annotated clinical variables, the 34 genes showed a strong positive correlation with therapeutic response (**Fig 2B-D**), as determined by endoscopic and histologic evidence of mucosal healing—an objective and stringent endpoint commonly used in IBD clinical trials^45^.

**Figure 2.**
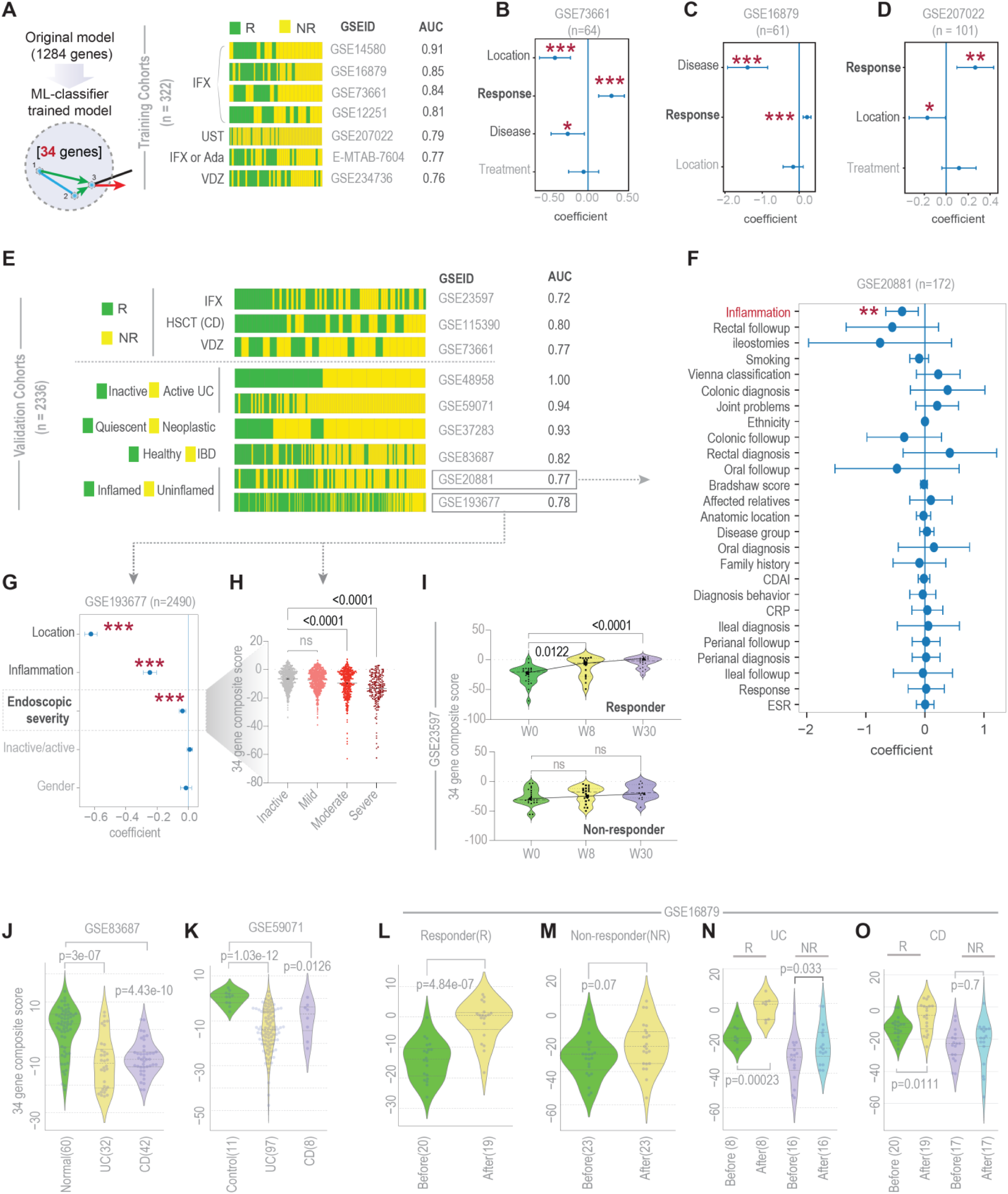
Identification of a molecular signature of therapeutic remission and derivation of a therapeutic goal. **A**. Bar plots display the ability of the 34-gene signature to classify responder vs non-responder in the training datasets. Performance is measured by computing ROC-AUC. The rank order of different samples on the bar plot is based on the composite scores of 34 genes, arranged from left (high score) to right (low score). HSCT, Hematopoietic stem cell transplantation. **B-D**. Multivariate analyses on the training cohorts (in B) that reported >/= 3 variables (including ‘response’ status) using the composite score of 34 genes as a linear combination of all variables. The coefficient of each variable (at the center) with 95% CI (as error bars) and the p-values are illustrated. The p-value for each tests the null hypothesis that the coefficient is equal to zero (no effect). **E**. Bar plots show the ability of the composite expression score of the 34 genes in rank-ordering IBD disease groups, as in B, in various discovery cohorts. The disease groups include: (*from top to bottom*) response status in 3 additional treatment cohorts (Infliximab, GSE23597; Vedolizumab, GSE73661; and hematopoietic stem cell therapy, GSE115390), disease activity, evidence for neoplastic progression, presence or absence of disease, and presence or absence inflammation. **F-G**. Multivariate analyses show the composite score of 34-gene signature as a linear combination of all other variables. Endoscopic severity in panel H is used as a continuous variable. The p-value for each term tests the null hypothesis that the coefficient is equal to zero (no effect). p-values are as follows: * = p ≤ 0.05, ** = p ≤ 0.01, and *** = p ≤ 0.001. **H**. Scatter plots display the composite score of 34 genes in samples annotated for varying endoscopic severity. The p values, calculated by Welch’s t-test are displayed. **I-O**. Violin plots display the composite score of the levels of expression the 34 genes in patients receiving biologics, categorized based on treatment duration (I), disease type (J-K), timing of biopsy (before vs after biologics; L-M) and response status by disease subtype (UC vs CD; N-O). p-values were calculated by Welch’s t-test. See also **Fig S2D-G** for functional enrichment analyses on the 34 gene signature. A catalog of all datasets and patient characteristics used in this work can be found in **Supplemental Datasheet 1**.

Validation studies confirmed that the 34-gene model reliably classified responders from non-responders regardless of therapeutic modality (biologics or cell-based therapies) or mechanism of action (anti-inflammatory, anti-trafficking, or regenerative) (**Fig 2E***-top*). Furthermore, the 34-gene signature of barrier dysfunction successfully stratified patient samples across additional validation cohorts: downregulation of this signature was a shared molecular feature among patients with active disease, inflammation, or neoplasia risk (**Fig 2E***-bottom*). We next tested the relevance of the 34-gene signature in complex, heterogeneous cohorts annotated with clinical outcome data. Multivariate analyses confirmed consistent inverse correlations between the 34-gene expression levels and both histologic inflammation (**Fig 2F**) and endoscopic severity (**Fig 2G-H**).

Composite expression score analysis of the 34-gene signature revealed that upregulation during treatment was a hallmark of therapeutic response (**Fig 2I**). This pattern was observed consistently in two cohorts and across disease subtypes, with suppressed expression in both ulcerative colitis (UC) and Crohn’s disease (CD) at baseline (**Fig 2J-K**). Post-treatment comparisons showed significantly greater upregulation among responders than non-responders (**Fig 2L-M**), and this trend was preserved across both disease types (**Fig 2N-O**).

Gene ontology analysis using multiple approaches showed significant enrichment of core bioenergetic processes among the 34 genes, including mitochondrial electron transport, cytochrome C oxidase activity, sulfur metabolism, and detoxification of hydrogen sulfide (H₂S) (**Fig S2D-G**).

Together, these findings indicate that the 34-gene signature reflects the gut’s underlying bioenergetic state—a key determinant of clinical response. Responders exhibited higher baseline expression and continued induction of this signature during treatment, whereas non-responders had lower baseline levels and/or failed to upregulate it over time. Most critically, this signature closely tracked endoscopic and histologic evidence of mucosal healing at treatment completion— a key endpoint in clinical trials. Thus, pharmacologic induction of the 34-gene signature emerges as a rational therapeutic goal.

### A framework of network-based rules for target prioritization

We next asked whether the 34 remission-associated genes were functionally connected—via BIRs—to genes encoding protein targets of drugs used in the training cohorts, such that their induction might be achieved through target perturbation. We compiled a comprehensive list of these drug target genes (**Supplemental Datasheet 2**). For each, we computed two key metrics: (i) Therapeutic index (**TI**); and (ii) Connectivity index (**CI**), which together informed a composite Likelihood of approval index (**LoAI**) for each drug (**Fig 3A**; see *Methods*). These calculations were based on the same high-quality IBD transcriptomic dataset used to build the original Boolean implication network model^16^, characterized by >90% read alignment and derived from full-thickness, surgically resected tissue from biologic-naïve patients^46^.

**Figure 3.**
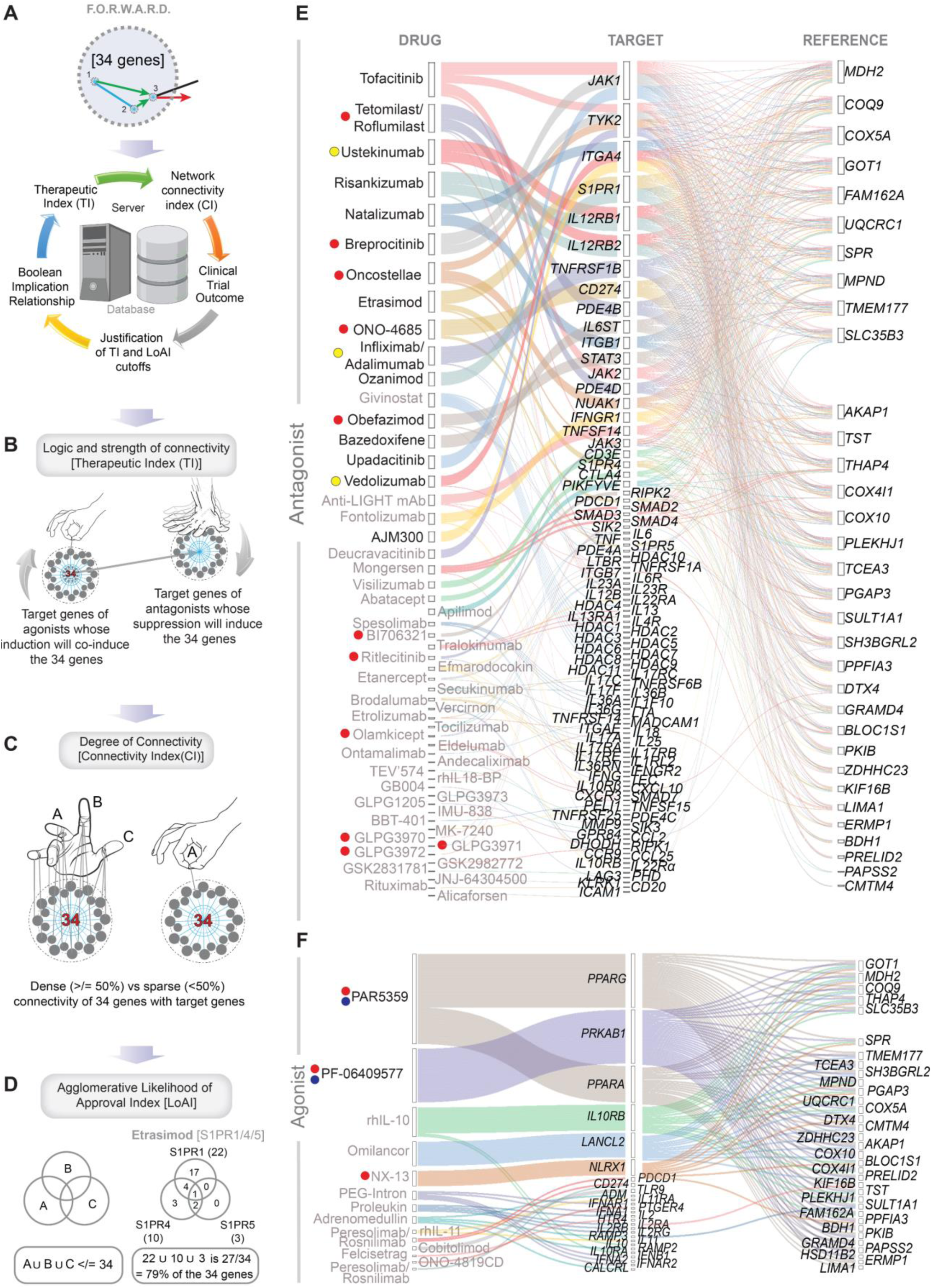
Connectivity of drug target gene(s) to the 34-gene molecular signature of therapeutic remission. **A.** Schematics illustrate the various statistical derivatives within the computational framework of F.O.R.W.A.R.D that enable objective scoring of the logic, strength and degree of connectivity of any drug’s target gene(s) with the 34 reference genes that track the likelihood of therapeutic remission. **B.** The logic and strength of connectivity is assessed by calculating ’therapeutic index,’ (see *Methods* and **Fig S1**). Drugs acting as *ago*nists are expected to co-induce the 34 reference genes via equivalent Boolean Implication Relationships (BIRs), ‘T equivalent R’ and ‘T high=> R high’ (*left*), whereas drugs acting as *anta*gonists are expected to induce the 34 reference genes via opposing BIRs, ‘T opposite R’ and ‘T low=> R high’ (*right*). **C.** Connectivity index (CI) is a number between 0 and 1, which reflects the fraction of the 34 reference genes to which target gene(s) relates to a TI <0.075. Connectivity is deemed as sparse vs dense using a >/= 50% as cutoff, based on the simple majority protocol. **D**. Schematic (*left*) showcases how the agglomerative likelihood of approval index (LoAI) for a generic drug is derived based on the union summation of its target gene’s connectivity with the 34-gene signature of remission. Schematic (*right*) cites the process of LoAI estimation for *Etrasimod*, which targets proteins encoded by 3 distinct genes. The CI for each target is indicated in parentheses. **E-F.** Alluvial plots for antagonists (E) and agonists (F) representing the connectivity stream between drug names (*left column*), major score-driving target gene (*middle column*) and the reference genes within the 34 gene signature of therapeutic remission (*right column*). A comprehensive list of target genes, TI, CI, and LoAI values, and their connectivity with reference genes are shown in **Supplemental datasheet 2**. *Black* and *grey* fonts indicate drugs with an LoAI of at least >/= 0.5 (i.e., achieved simple majority) or below (fails to reach majority), respectively. Red dots (in E-F) mark drugs in ongoing trials where outcomes are unknown currently. Yellow dots (E) mark the four FDA-approved drugs used for model training. Blue dots (F) mark promising preclinical targets identified using Boolean Implication network approach.

The Therapeutic index quantifies both the nature and statistical strength of BIRs between a query (target) gene and each of the 34 reference (R) genes (**Fig S1A-E**). TI indicates whether a target gene should be upregulated (via an agonist) or downregulated (via an antagonist) to induce the desired activation of the 34-gene signature (**Fig 3B**). The strength of these associations estimates the likelihood of achieving intended (inductive) or unintended (suppressive) effects on remission-associated gene expression.

The Connectivity index measures the fraction of the 34 reference genes that are connected to a given target gene using a stringent TI threshold of <0.075. By a simple majority rule, targets with dense connectivity (<50% of 34 reference genes) are prioritized over those with sparse connectivity (>/= 50% of 34 reference genes) (**Fig 3C**). This majority rule is supported by its effectiveness in achieving consensus and controlling behavior in complex, fully connected networks, particularly when information transfer is noisy or incomplete^47^.

Because many drugs act on multiple targets, we calculated a drug-specific LoAI as the cumulative, non-redundant union of CI scores across all target genes of each drug—ensuring that each of the 34 reference genes is counted only once (see *example*; **Fig 3D**).

For the four drugs in the training cohorts that advanced to phase 3 and received regulatory approval (**Fig 3E**-*yellow dots*), primary antagonistic targets showed >50% connectivity with the 34 reference genes (**Fig 3E**-top, *black*). By contrast, the failed drug programs—such as those targeting *SMAD7*, *MAdCAM-1*, *ITGB7* and *IL36*---demonstrated sparse connectivity (**Fig 3E**-bottom, *grey*). The LoAI also captured cases of multi-target drugs where contradictory effects on reference genes canceled each other out, as in the failed pan-HDAC inhibitor givinostat (**Fig S3A-B**; NCT00792740). Full CI values for each drug-target pair are in **Supplemental Datasheet 2**.

This connectivity framework also grouped current investigational targets (**Fig 3E**-red dots) with either successful or failed historical programs, providing predictive insight. Given the lack of phase 3 success for agonists, we analyzed all previously failed agonist targets alongside current candidates (**Fig 3F**), including two newly identified preclinical assets that have emerged from the Boolean transcriptomics: the PRKAB1-agonist PF-06409577^16^ and the balanced PPARα/γ-dual agonist PAR5359 ^17^. Notably, none of the historical agonists, all of which had failed clinical trials surpassed the 50% connectivity threshold.

These findings suggest that drugs with successful phase 3 outcomes tend to show stronger and more extensive connectivity—across logical, statistical, and functional dimensions— with the 34-gene remission signature. Because this network-based framework leverages real-world clinical outcomes to assess target-drug compatibility, we refer to it as *F.O.R.W.A.R.D*: Framework for Outcome-based Research and Drug Development.

### Benchmarking F.O.R.W.A.R.D against other network-based approaches

We next evaluated how F.O.R.W.A.R.D. compares with other established target identification strategies (**Fig 4A-B**). For example, Differential expression analysis (DEA), widely regarded as the gold standard, underpins several generative-AI approaches for target discovery, including PandaOmics by Insilico Medicine^31^. Similarly, a Bayesian method that integrates genomic, transcriptomic, and clinical data from IBD patients has also been used to construct functional genomic networks, identifying 133 key driver genes (KDGs^46^), of which a top-ranked 12-gene subset was validated as key regulator of IBD pathophysiology. We also benchmarked against a 74-gene machine learning (ML) signature derived using a bagged trees algorithm as IBD drivers^48^. Gene overlaps across the four signatures were limited (**Fig 4A**). To compare predictive value, we performed univariate analyses on clinical trial datasets to assess each signature’s correlation with treatment response, the primary outcome of interest (**Fig 4C**). F.O.R.W.A.R.D’s 34-gene signature demonstrated consistent, significant positive correlation with response in 8 of 9 cohorts. The 133-gene Bayesian KDGs reached significance in 7/9 cohorts but showed inconsistency in correlation with response (negatively correlated in all cases, except in GSE76331; **Fig 4C**-*3^rd^ plot from the left*). The 12-gene Bayesian subset, DEA, and ML signatures showed weaker and inconsistent performance. Notably, the 1284-gene Boolean network superset from which F.O.R.W.A.R.D. was derived outperformed conventional approaches (significant in 7/9 cohorts) but still underperformed relative to its 34-gene outcome-trained subset.

**Figure 4.**
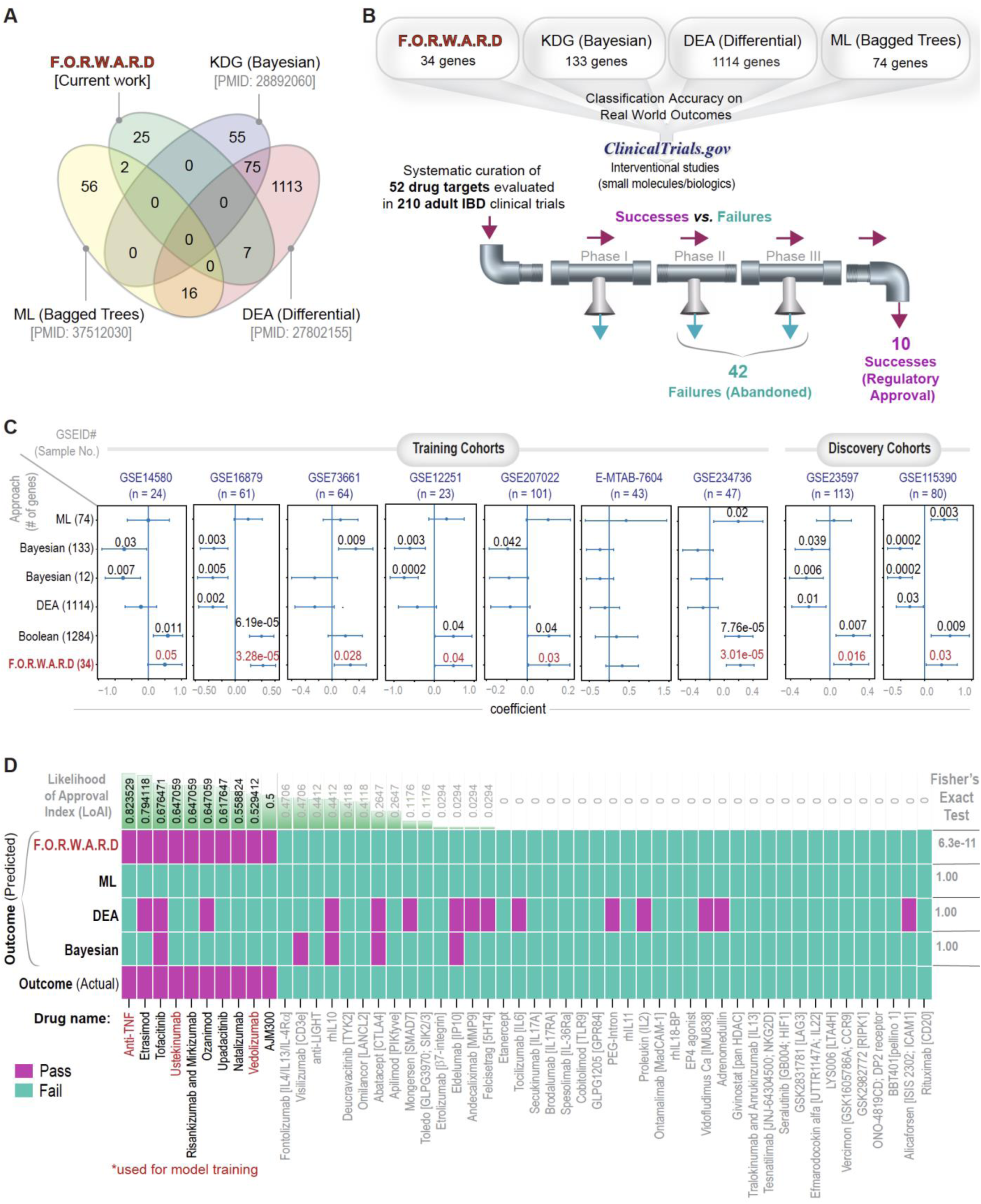
Predictive accuracy of F.O.R.W.A.R.D on drug development programs with known outcomes. **A**. Venn diagram illustrating the overlap of genes identified by various methods: the Bayesian KDG algorithm^46^, Differential expression analysis (DEA) between healthy individuals and IBD patients^60^, the bagged tree algorithm ^96^, and the 34 genes used in F.O.R.W.A.R.D (current work). ML, machine learning. **B** Schematic illustrates the process by which drug programs were chosen for their inclusion in benchmarking studies (see details in **Fig S4**). A catalog of all drug programs included in this analysis can be found in **Supplemental Datasheet 2** and gene signatures in **Supplemental Datasheet 3**. A flowchart showing steps and criteria of systematic curation of drug targets can be found at Supplemental Figure S3. **D**. Univariate analyses presenting the composite scores of various gene sets in A, and their related signatures, i.e., the 12 top-ranked KDGs that is a subset of the 133 Bayesian KDGs^46^, and the superset of 1284 genes, from which the 34 genes were derived for the F.O.R.W.A.R.D algorithm (**Fig 2A**, current work). Results are displayed as a linear combination of all the composite scores with the response status of patients across all training and validation cohorts. The coefficient of each variable (centered) is displayed with 95% confidence intervals (error bars) and the numbers display p-values. The p-value for each tests the null hypothesis that the coefficient is equal to zero (no effect). **E.** A binary matrix heatmap shows the pass (magenta) and fail (teal) prediction on 52 targets representing 210 clinical trials based on F.O.R.W.A.R.D and 3 conventional methods that are currently used to prioritize target genes. The three drugs that were used for training F.O.R.W.A.R.D are highlighted in red. The actual outcome is annotated in the bottom row. The Likelihood of approval (LoAI) score derived from the F.O.R.W.A.R.D framework is displayed across the top row. Each drug’s target genes, and their TI, CI and LoAI are presented in **Supplemental datasheet 2.**

To benchmark clinical performance, we curated a comprehensive list of successful targets from FDA.gov and international approvals [e.g., Filgotinib in Japan, Great Britain and the EU; AJM300 (Carogra®)^49^ in Japan], and a list of failed targets from ClinicalTrials.gov, the EMA Clinical Trials Register, and industry press releases. To focus on efficacy, early-stage programs terminated due to safety concerns were excluded, yielding 52 targets tested in 210 trials involving adults with IBD (see inclusion/exclusion criteria in **Fig S4**).

F.O.R.W.A.R.D. significantly outperformed all other models in distinguishing successful (LoAI ≥ 0.5) from failed (LoAI < 0.5) targets (Fisher’s exact test: 6.3e-11; **Fig 4D**). Notably, Mongersen (GED-301), which met its primary endpoint in phase 2 but failed in phase 3^49^, scored 0 in the FORWARD model (**Fig 4D**). Etrolizumab, an anti-β7 integrin antibody with positive phase 2 results^50^ and five lackluster phase 3 trials^38,51–55^, also failed to gain approval and scored 0.029 (**Fig 4D**). Cobitolimod, a TLR9 agonist that succeeded in phase 2^56^ but failed in phase 3 (NCT04985968), scored 0 (**Fig 4D**). Ontamalimab, which targets MadCAM-1 and was tested in six lengthy phase 3 trials^57^, was terminated and also scored 0 (**Fig 4D**).

These findings highlight that F.O.R.W.A.R.D., based on a 34-gene remission signature, objectively and effectively predicts regulatory success across therapeutic modalities and mechanisms, offering a method for target prioritization grounded in clinical outcome data.

### An epithelial 34-gene signature tracks future endoscopic remission

To investigate the cellular origin of the 34-gene remission signature, we analyzed two independent single-cell transcriptomic datasets: one from colonic biopsies of healthy individuals and IBD patients (GSE214695), and another from a longitudinal cohort of biologic-naïve IBD patients undergoing anti-TNFα therapy with follow-up for therapeutic response (GSE282122). Using AUC-based classification, we assessed each gene’s ability to discriminate between disease and response status across cell types. When it came to classifying disease, the 34-gene signature was predominantly upregulated in epithelial lineages—particularly tuft cells, transit-amplifying (TA) cells, goblet cells, and colonocytes—in healthy individuals, with high classification accuracy (**Fig 5A**). When it came to classifying future therapeutic response to anti-TNFα therapy, expression of the signature was elevated in tuft and TA cells of responders (**Fig 5B**). TA cells emerged as the most consistent site of strong expression across both healthy and treatment-responsive individuals, suggesting specificity to a pool of rapidly proliferating progenitors that arise from intestinal stem cells and are responsible for generating the bulk of the crypt epithelial cells. This pattern suggests that the 34-gene signature might be epithelial-specific. To confirm this, we computed the composite 34-gene score in TA cells across both datasets. Scores were significantly higher in healthy versus IBD patients (**Fig 5C***-top*) and in responders versus non-responders (**Fig 5C***-bottom*). This epithelial association was further validated using laser-capture microdissected colonic epithelium (TA accounts for ∼25% of the total crypt cell population^58^) from an independent cohort, which showed similar upregulation of the 34-gene signature (**Fig 5D**).

**Figure 5.**
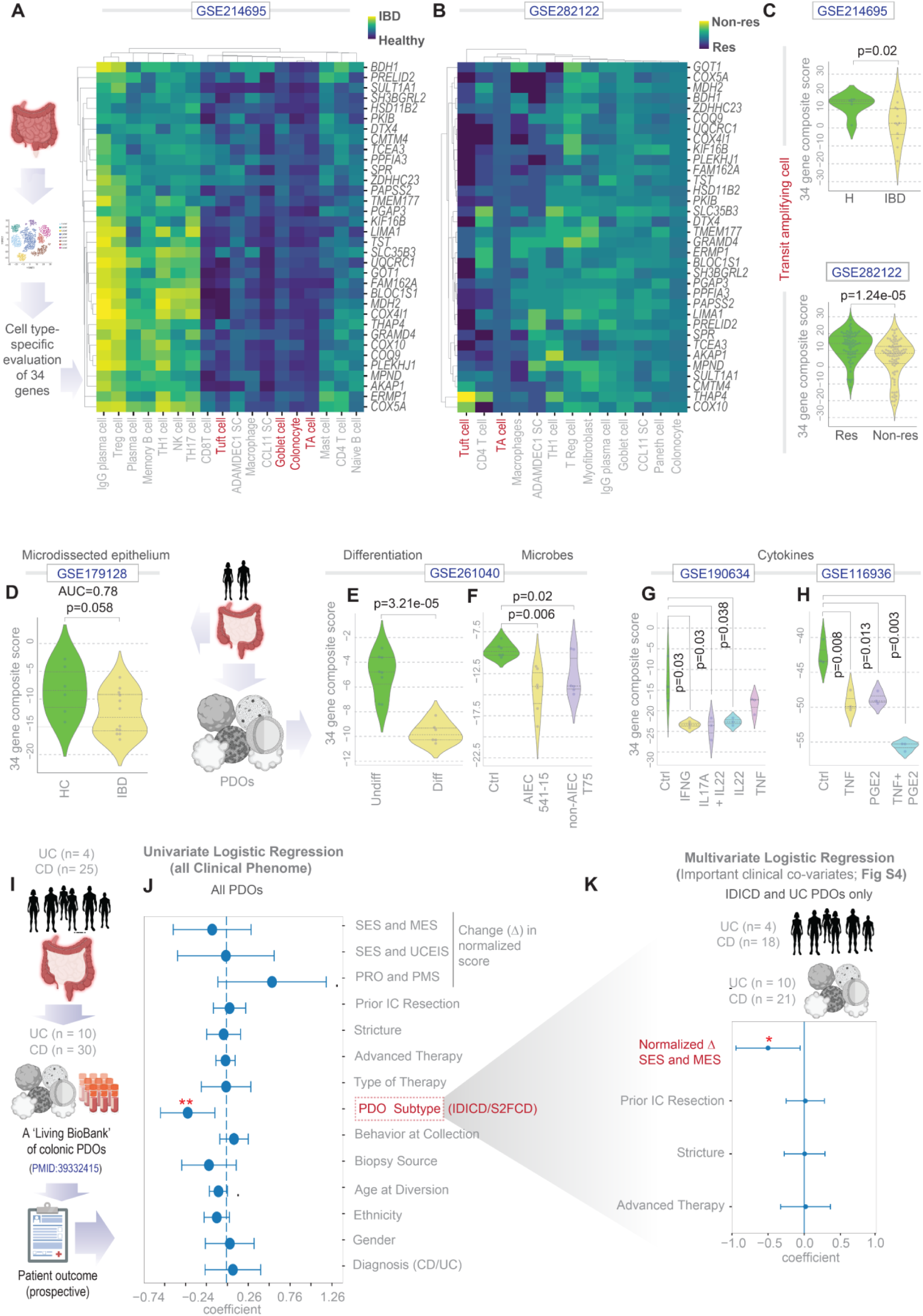
Validation of the predictive potential of an epithelium-specific 34-gene molecular signature. **A-B**. Workflow for assessing cell-type specific expression of 34 genes in single cell transcriptomic datasets from colons (A-*left*). Heatmaps show classification accuracy between healthy individuals and IBD patients (A-right) and between responder and non-responders undergoing anti-TNFα therapy (B). TA, transit amplifying. **C**. Violin plots display the composite expression score of the 34 genes only in TA cells from the datasets analyzed in A and B. **D-H**. Violin plots display the composite expression score of the 34 genes in micro-dissected colon epithelium isolated from healthy and IBD patients (D), PDOs derived from differentiated and undifferentiated monolayers (E), and monolayers treated with adherent and non-adherent *E. coli* (F) or with various combinations of cytokines (G-H). **I**. Workflow of prospective study design using PDOs as *ex vivo* platforms for assessing the relationship between epithelium-specific expression of the 34-gene signature and disease remission. **J**. Univariate model of the composite score of the 34-gene signature as a base variable, tested against each clinical covariate collected over a 5-year prospective follow-up in a previously curated, genotyped-phenotyped PDO cohort^21^. SES, Simple endoscopic score for Crohn’s disease; MES, Mayo endoscopic score; UCEIS, Ulcerative Colitis Endoscopic Index of Severity; PRO, patient-reported outcome; PMS, partial Mayo Score. The three variables on the top of the list were scaled across molecular subtypes. See also **Fig S5** for univariate model on exclusively the IDICD/UC population. **K**. Multivariate analysis depicting the composite score of the 34-gene signature as a linear combination of clinically important covariates within IDICD/UC group. For both J and K, the coefficient of each variable (at the center) with 95% CI (as error bars) and the p values are illustrated. The p-value for each term tests the null hypothesis that the coefficient is equal to zero (no effect). p ≤ 0.1; *p ≤ 0.05; **p ≤ 0. 01. Bold = significant co-variates.

To explore environmental and regulatory influences on signature expression, we leveraged public datasets from patient-derived organoids (PDOs), which serve as dynamic platforms to model TA cell’s contribution to crypt regeneration in vitro^58^. The composite 34-gene score was significantly lower in differentiated monolayers (which are depleted of TA cells) compared to undifferentiated PDOs (**Fig 5E**). This downregulation was further enhanced by microbial exposure—particularly with adherent *E. coli* strains, commonly enriched in IBD microbiota (**Fig 5F**). Additionally, treatment of differentiated PDOs with pro-inflammatory cytokines (TNFα, IFNγ, IL-17A, IL-22, PGE2), alone or in combination, consistently suppressed 34-gene composite scores (**Fig 5G–H**). These findings reinforce that inflammatory stress and microbial colonization, key features of IBD pathology, repress this epithelial-specific remission signature.

To assess the prognostic potential of this signature, we analyzed a living biobank of PDOs from the UC San Diego IBD Biobank, with prospectively collected clinical data over 4–5 years (**Fig 5I**; metadata in **Supplementary Datasheet 4)**. The PDOs included samples from 25 Crohn’s disease (CD) patients, stratified into two molecular subtypes—Immune-Deficient Infectious CD (IDICD) and Stress- and Senescence-Induced Fibrostenotic CD (SF2CD)—along with 4 UC patients^21^. Based on prior PCA analysis^21^, UC-derived PDOs were grouped with IDICD-like epithelia.

Univariate regression using the composite 34-gene score identified molecular subtype as the only significant explanatory variable (p ≤ 0.01; **Fig. 5J**). Specifically, the 34-gene predictor significantly correlated with the IDICD/UC PDO subtype. To evaluate clinical relevance, we performed univariate (**Fig S5**) and multivariate regression (**Fig 5K**) within the IDICD/UC subtype using the 34-gene composite score as a predictor. We retained all clinical covariates for the univariate model (**Fig S5**) and selected only the most relevant covariates [e.g., endoscopic evidence of remission during follow-up] for the multivariate model (**Fig 5K**). In both models, lower 34-gene scores were significantly associated with persistent or recurrent disease activity, as measured by normalized Simple Endoscopic Score (SES) for CD and Mayo Endoscopic Score (MES) for UC (p ≤ 0.01; **Fig. 5K**). The consistent significance across both models suggests the 34-gene score is an independent predictor, unlikely to be confounded by other clinical variables.

Together, these results support the 34-gene remission signature as (i) epithelial in origin, (ii) suppressed by inflammation and microbial stress, and (iii) prognostic of disease activity, positioning it as a clinically actionable biomarker in IBD.

### F.O.R.W.A.R.D enables dynamic target prioritization and in silico trial design

F.O.R.W.A.R.D can be integrated at multiple stages of therapeutic development. In early clinical phases, it supports the vetting and prioritization of individual targets or gene combinations (**Fig 6A***-left*), or broader screening across entire target classes (e.g., kinome, GPCRome). In later phases, F.O.R.W.A.R.D can be trained on emerging outcomes to enable iterative refinement (**Fig 6A***-right*). Beyond target discovery, we tested its utility in guiding clinical trial design and cohort selection through *in silico* trials.

**Figure 6.**
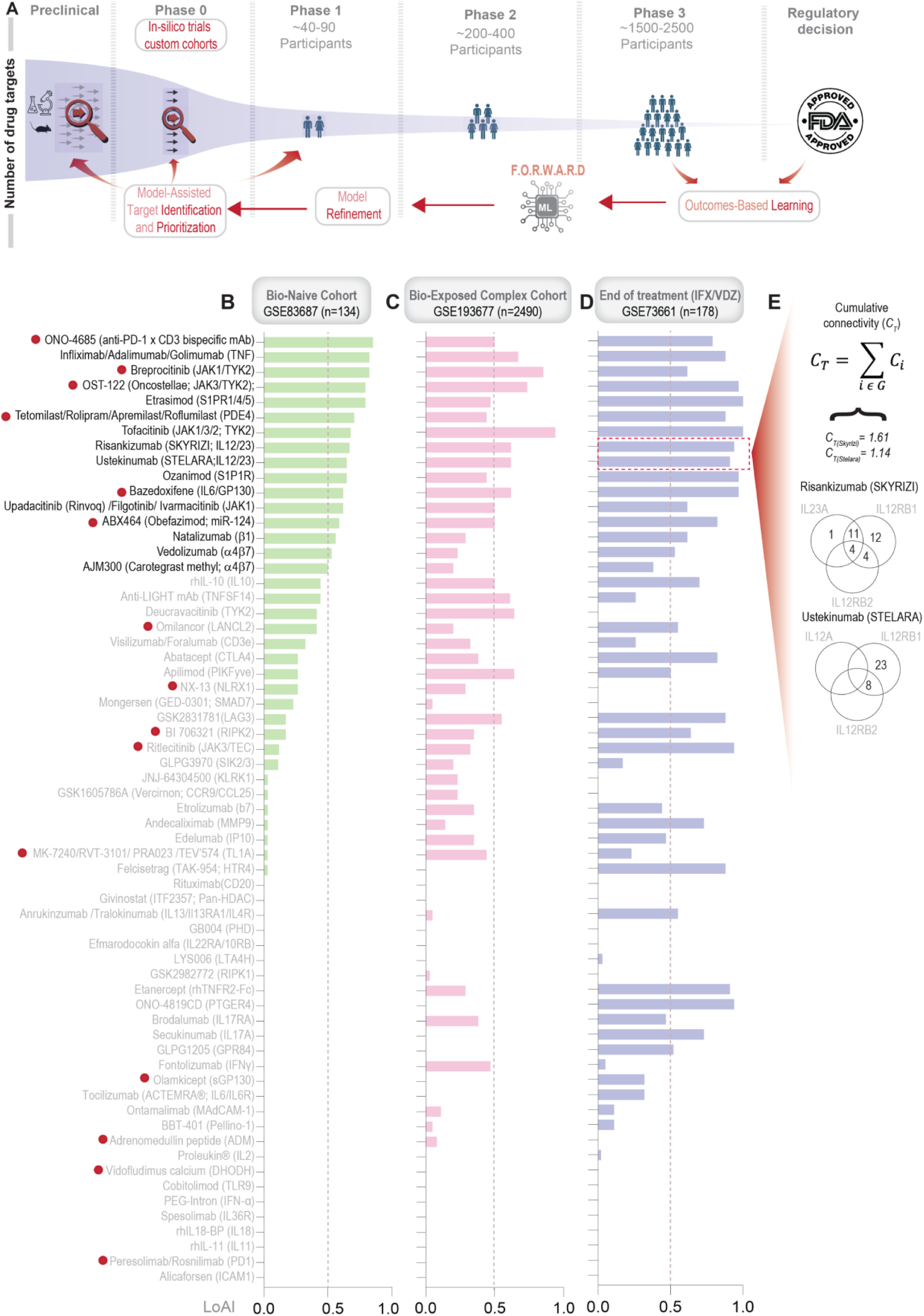
Integration of F.O.R.W.A.R.D into the current drug discovery process through in silico trials. **A.** Schematic shows how the F.O.R.W.A.R.D framework could be seamlessly integrated into both the early and late stages of the drug development process (*red* arrows) and the potential impacts of such integration (*red text* in boxes). **B-D**. Bar plots show the likelihood of approval index (LoAI) of drugs and their major target genes (listed in parenthesis) in three custom cohorts reflecting important clinically directive characteristics considered during trial design and recruitment. Red dots next to a drug’s name implies that these are ongoing trials. IFX, infliximab; VDZ, vedolizumab. **E.** Cumulative connectivity, which is an estimate of total overlapping connectivity of multi-target drugs with the 34-gene signature of response, is shown for risankizumab, mirikizumab and ustekinumab, as determined in a cohort of patients exposed to anti-TNF or vedolizumab.

While the initial IBD network and 1284-gene barrier signature were validated using ∼2700 diverse patient samples (**Fig 1***-step 1*), network connectivity between drug targets and the 34- gene remission signature was evaluated only in a biologic-naïve cohort (**Fig 3E-F**). To assess how prior biologic exposure might alter predicted efficacy, we calculated LoAI scores across three *in silico* trials: (i) A biologic-naïve cohort^16^ (original Boolean Implication network; **Fig 6B**); (ii) A real-world-like complex cohort^59^ (∼75% late-stage biologic-exposed; **Fig 6C**); and finally, (iii) A cohort with acute biologic exposure^60^ (4-6 weeks after a single dose of infliximab or 6, 12 and 52 weeks post-treatment with vedolizumab from two phase 3 trials, GEMINI I and LTS). Comparative LoAI analysis revealed striking substantial shifts in predicted drug efficacy across these cohorts (**Fig 6B-D**). Consistent with the concept of dynamic network rewiring in chronic disease^61^, targets like JAK, TYK2, PIKfyve, and LIGHT gained predicted efficacy in the complex setting, whereas α4β7 (vedolizumab), S1PR1 (ozanimod), and S1PR1/4/5 (etrasimod) lost efficacy. IL12/23 inhibitors—ustekinumab, risankizumab, and mirikizumab—retained efficacy across all settings. Existing evidence suggests that at least some of these predictions are true (see *Discussion*). Strikingly, several previously failed drugs—e.g., CTLA4, IL17A, MMP9, and IL10 antagonists— were predicted to be effective if re-trialed following acute biologic exposure (**Fig 6D**). While these predictions remain to be clinically validated, they suggest a rationale for sequential therapy trials. Conversely, the pan-HDAC inhibitor givinostat consistently underperformed across all cohorts and was predicted to exacerbate disease in the most complex setting (**Fig S3B-D**).

To explore predictive resolution, we tested whether F.O.R.W.A.R.D could differentiate among competing therapeutics. All three IL12/23 inhibitors exceeded the LoAI threshold in the naïve cohort (**Fig 6B**), but risankizumab and mirikizumab showed higher scores and stronger cumulative connectivity (C_T_) to the reference gene set in the bio-exposed cohort (**Fig 6D-E**), consistent with their superior performance in both real-world and direct head-to-head trials (e.g., SEQUENCE^62^ trial).

Together, these findings highlight F.O.R.W.A.R.D’s potential to guide target prioritization, inform rational trial design, identify opportunities for re-trials, and predict relative efficacy across therapeutic candidates.

## Discussion

The major innovation we present here is a computational framework designed to enhance precision in target discovery. Below, we outline five key innovations and their implications:

### Molecular Goal-Driven Targeting Through a 34-Gene Signature

F.O.R.W.A.R.D prioritizes targets based on a well-defined therapeutic goal: upregulating a validated 34-gene remission signature by perturbing proximal network nodes with drugs. This approach reduces bias often introduced by conventional enrichment analyses^31,32^ and enhances the discovery of novel, biologically grounded targets. The 34-gene signature—strongly linked to mitochondrial function and bioenergetics—emerged as a robust marker of gut barrier integrity, remission, and response. Gene ontology enrichment implicates this signature in mitochondrial bioenergetic pathways—including biogenesis, respiration, redox balance, and quality control— hallmarks of epithelial dysfunction in IBD^63–66^. Notably, none of the 34 genes were linked to IBD risk or severity in GWAS studies, suggesting a potential role for epigenetic regulation, supported by emerging literature on epigenetic control of metabolic remodeling in IBD^67–72^.

### Predictive Performance Exceeds Conventional Methods

Compared to standard approaches—differential expression analysis (DEA), artificial neural networks (ANNs), and Bayesian inference—F.O.R.W.A.R.D demonstrated superior ability to identify clinically successful targets. This is likely because the BIR framework captures both symmetric and asymmetric gene-gene relationships, allowing it to recognize biologically meaningful associations missed by correlation-based traditional methods. The BIR framework is also likely to have mitigated the risk of overfitting because they capture asymmetric gene expression relationships reflecting stable, biologically grounded logic rather than spurious correlations as demonstrated in our previous works^16,24^. Unlike conventional statistical methods, BIRs impose structural constraints that reduce model complexity and enhance generalizability across diverse datasets. We believe that this combination of biologically constrained logic and rigorous model selection yielded a robust and interpretable framework. The strong translational potential of the framework is evident in its ability to flag numerous historical failures with high precision (see **Fig 4D**), indicating that objective LoAI scoring could have supported early termination. Its translational potential is also validated prospectively in PDOs (see **Fig 5J-K**; **Fig S5**). Finally, the platform also offers predictive insights into ongoing trials: TL1A antagonism, evaluated in phase 2 studies (DIONE, TUSCANY^73^ and TUSCANY II^74^), is predicted to succeed phase 3 only if the trial were to be performed in biomarker-enriched populations. These results highlight F.O.R.W.A.R.D’s exactness and utility in derisking drug development through early-stage prioritization or discontinuation.

### Predictive Potential of Epithelial-Specific Expression of the 34-Gene Signature

Single-cell and microdissected datasets revealed that the 34-gene signature is highly enriched in epithelial tuft and TA cells and is preserved in healthy tissue and in therapy responders. This compartment-specific expression pattern was consistent across multiple platforms and aligns with the known role of epithelial bioenergetics in mucosal healing. Using PDOs as ex vivo models of the gut epithelium, we demonstrated that reduction in TAs (during differentiation) or epithelial injury (exposure to microbes or inflammatory cytokines) downregulate this signature, consistent with the dysbiotic and inflamed mucosal environment in IBD. Further, PDOs from subjects with the molecular subtype that captures both microbial dysbiosis and inflammation, i.e., IDICD/UC subtype that had lower 34-gene expression, were less likely to achieve or stay in remission. These findings establish the 34-gene signature as a cell-type-specific, clinically measurable marker of disease activity and remission potential, with utility in patient stratification and outcome prediction.

### Integration of F.O.R.W.A.R.D into In Silico Trials and Trial Design

We demonstrate the utility of F.O.R.W.A.R.D as a decision-support tool at critical stages of drug development, including virtual phase 0 trials for target assessment and cohort-specific predictions. As clinical trial designs grow increasingly stringent^75^, enrolling sufficient biologic-naïve patients has become a major bottleneck^76^, underscoring the need for innovative trial strategies^77^. F.O.R.W.A.R.D. addresses this by simulating therapeutic outcomes across customized virtual cohorts—stratified by disease severity, subtype, stage, and prior biologic exposure—offering actionable insights to inform trial design. Importantly, this cohort-specific predictive capability is not a limitation but a strength. In chronic disease, network rewiring^61^ shifts target relevance over time, and F.O.R.W.A.R.D’s predictions reflect this underlying biological logic—capturing meaningful dynamics rather than relying on static or spurious associations. While prospective validation is pending, several predictions align with real-world data. instance, tofacitinib has demonstrated greater effectiveness than vedolizumab in anti-TNF-exposed patients^78^, whereas IL-12/23 inhibitors maintain consistent efficacy across diverse clinical contexts^79,80^. Conversely, vedolizumab shows reduced efficacy in patients with prior anti-TNF exposure or more complex disease^81,82^. Findings also suggest that F.O.R.W.A.R.D can uncover therapeutic opportunities in previously failed programs—particularly when re-evaluated in specific patient subgroups or treatment sequences. As an example, IL-12/23 antagonists that pass F.O.R.W.A.R.D’s LoAI criteria can be further prioritized using cumulative connectivity (CT) analysis to predict comparative performance in head-to-head trials (see **Fig 6E**). By identifying sub-cohort–dependent efficacy patterns, F.O.R.W.A.R.D offers a rational pathway to repurpose underperforming assets and optimize clinical trial design.

### Continuous Learning from Trial Outcomes Can Enhance Predictive Accuracy

F.O.R.W.A.R.D is designed to evolve through iterative training on new datasets, including both failed and successful clinical programs. This refinement can optimize the 34-gene remission signature and recalibrate TI thresholds over time without modifying the CI majority rule. Although existing “learning” models for drug development exist^83^, they often rely on conventional DEA and enrichment methods ^31,32^; prone to annotation bias^32,84–90^, i.e., skewed towards richly annotated genes instead of those with the strongest molecular data^84^. F.O.R.W.A.R.D avoids this pitfall by leveraging direct molecular associations and Boolean logic. A recent example is the identification of TINK (TRAF2- and NCK-interacting kinase) as a fibrosis target using an enrichment-driven AI model; its lead candidate, INS018_055 is being tested in idiopathic pulmonary fibrosis (IPF)^91^. The program is in the early stages of phase 2, and outcomes have yet to emerge.

## Limitations

While F.O.R.W.A.R.D. can prioritize high-efficacy targets, it cannot account for druggability (e.g., potency) or toxicity—two critical factors in clinical success. For example, it does not explain why among two IL-12(p40)/23 antagonists, ustekinumab succeeded while briakinumab failed due to adverse effects^92^. While unable to model toxicity directly, F.O.R.W.A.R.D. can approximate off-target effects using strategies similar to those applied to pan-HDAC inhibitors. The platform also cannot capture efficacy gains derived from non-transcriptomic companion biomarkers [e.g., in TLA1-centric trials], unless the biomarker is transcriptome-based and can be simulated *in silico*.

Because F.O.R.W.A.R.D. relies on bidirectional influence relationships (BIRs) that infer regulatory mechanisms^15,22^, experimental validation remains essential, particularly where agonist/antagonist activity yields unexpected transcriptional outcomes. This is critical for targets where receptor activity diverges from transcript abundance, often in a dose-dependent manner^93,94^, as seen in the terminated Fingolimod (NCT01375179), and Ozanimod (YELLOWSTONE^95^) IBD trials. In such cases, F.O.R.W.A.R.D. supports revisiting therapeutic strategies at alternative doses to achieve molecular efficacy. Finally, our *in silico* simulations show F.O.R.W.A.R.D. requires recalibration of efficacy predictions within context-specific networks tailored to biologic-naïve or exposed populations—highlighting both a constraint in generalizability and a core strength in adaptability.

## Conclusions

F.O.R.W.A.R.D. introduces a data-driven approach to target selection in drug development that integrates real-world clinical insights with *in silico* network biology. Unlike deep learning “black-box” models, F.O.R.W.A.R.D. retains interpretability, ensuring outputs are transparent, actionable (humans-in-the-loop), and biologically grounded. Its modular and reproducible architecture supports seamless integration into existing discovery pipelines, offering immediate utility for enhancing precision and reducing risk in early-phase drug development. By learning from past failures, F.O.R.W.A.R.D. converts hindsight into foresight—enabling early identification of likely failures and re-optimizing misaligned programs for success. In doing so, it provides a powerful tool for reducing uncertainty and improving the strategic decision-making that underpins therapeutic innovation.

## Acknowledgments

No editing services or AI-technologies were used in drafting any part of this manuscript.

## Author contributions (CRediT)

Conceptualization: SaS, PG

Methodology & Resources: SaS, PG

Formal Analysis, Validation, Software: SaS

Consultation and domain expertise: BB, SiS.

Data Curation, Investigation & Visualization: SaS, EM, MM, PG

Funding acquisition: PG

Project administration: PG Supervision: PG

Writing – original draft: SaS, PG

Writing – review & editing: SaS, EM, MM, SiS, BSB, PG

## Data and materials availability

All data are available in the main text or the supplementary materials. Newly generated transcriptomic datasets reported in this paper involve re-analysis of GSE192819 which has been deposited in NCBI’s Gene Expression Omnibus. The new GSE ID assignment is GSE294760. All the publicly available datasets that were used in this work are listed in **Supplemental Dataset 1**.

## Supplementary Materials

Figs. S1 to S5

References (*1-96*)

Datasheet (Excel) S1 to S4

