## Supplementary Online Materials for "F.O.R.W.A.R.D: A Data-Driven Framework for Network-Based Target Prioritization in Drug Discovery"

**The PDF file includes:**

Supplementary Figures. S1 to S5 and legends
Supplementary Table 1
Details of Supplemental Datasheet 1 to 4
References

**Other Supplementary Materials for this manuscript include the following:**

Supplemental Datasheets 1 to 4

### Supplementary figures and legends

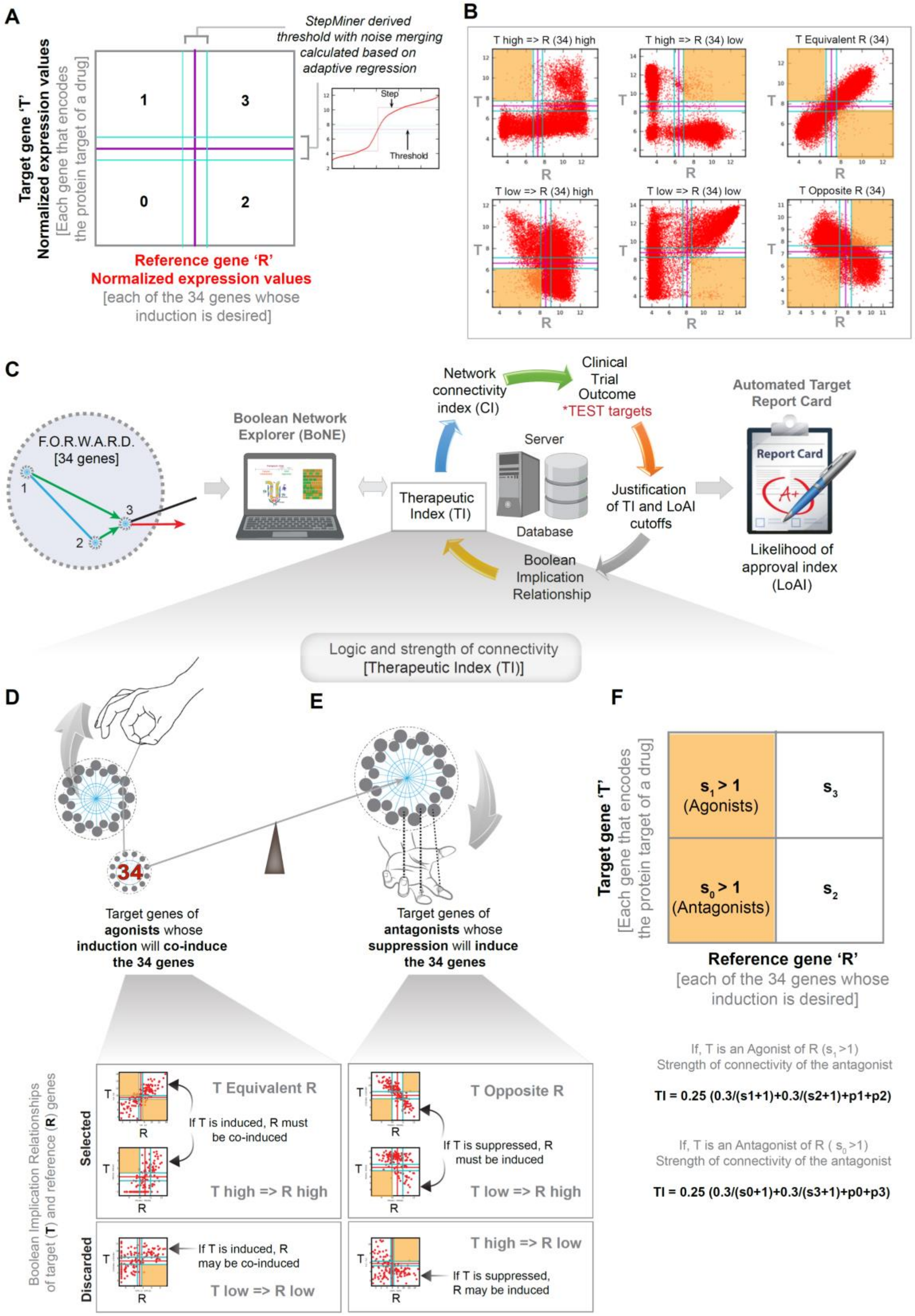


**Figure S1.** ***Boolean implication statistics and the derivation of the therapeutic index.***

**A.** Schematic displays the key principles behind the generation of scatter plot using normalized expression profile of any drug's target gene 'T' and any of the 34 reference 'R' genes. StepMiner-derived threshold and noise margins are represented using purple (threshold) and blue (noise margin) lines. **B.** Six possible Boolean implication relationships between any drug's target gene 'T' and any of the 34 reference 'R' genes. Sparse quadrants, computed using Boolean implication statistics, are shaded yellow. **C.** Schematic of computational framework involved in the calculation of the Likelihood of approval index (LoAI) of a drug based on the Therapeutic index (TI) and Connectivity index (CI) of its target gene (T) with the set of 34 reference (R) genes. **D-E.** Schematic representation of selected and discarded Boolean implication relationships between target genes of agonist (D) or target genes of antagonist (E) and 34 reference genes. **F.** Schematic showing sparse quadrant to be considered while calculating TI between agonist and antagonist drug target and expression of therapeutic index based on Boolean implication statistics (*S* and *p;* see Methods).

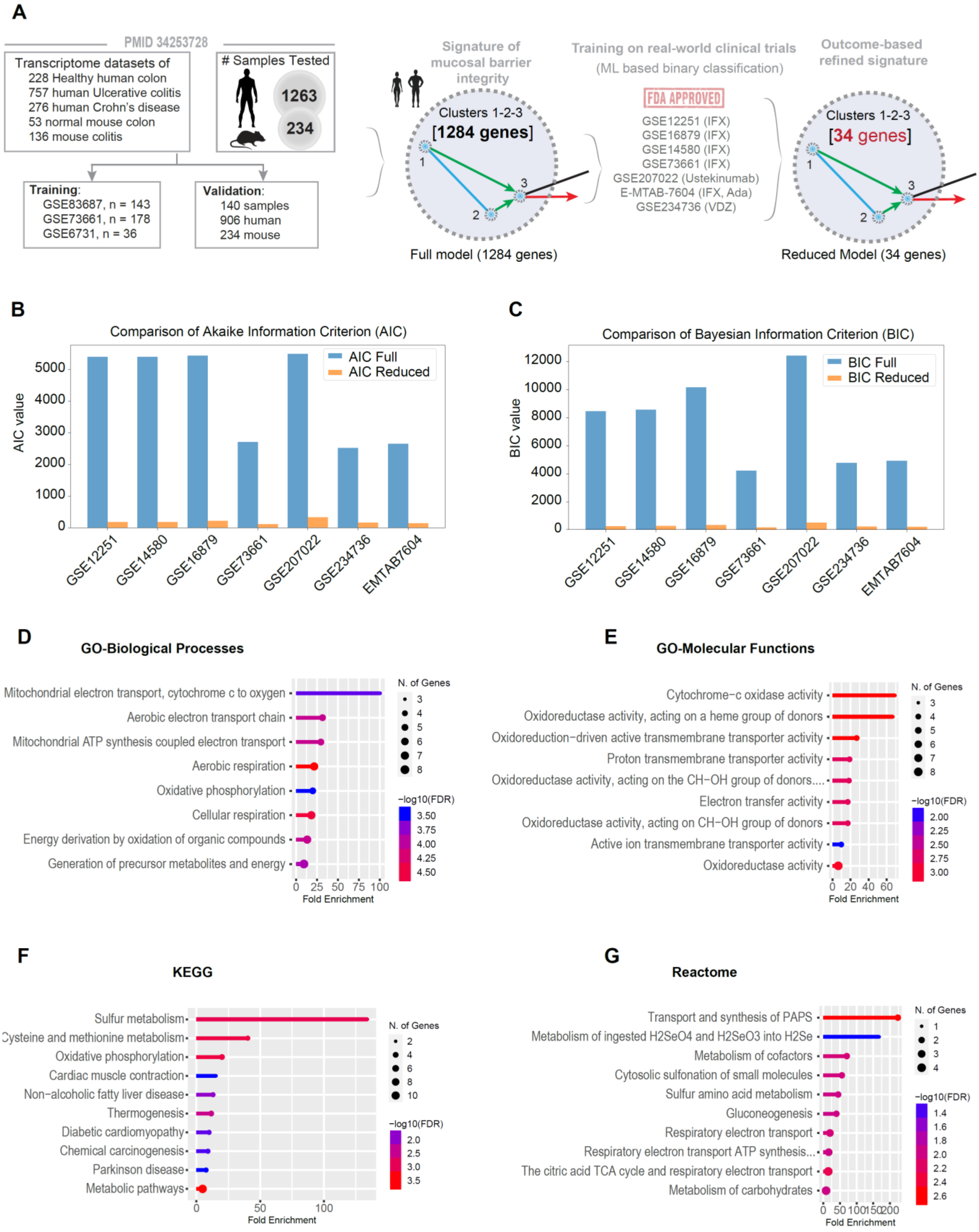


**Figure S2.** ***Model training and functional enrichment analyses.***

**A**. The computational workflow and the number of unique IBD samples that led to the identification of a 34-gene molecular signature of therapeutic remission. *Left*: Full model of 1284 genes (in gene clusters# 1, 2, and 3 of a previously validated Boolean IBD network[1]) that were identified using machine learning (ML) as the invariant signature of mucosal barrier integrity in all samples across species. *Middle*: The signature of barrier integrity (reduced model) is subsequently trained using ML-based classifier on pre-treatment samples in 7 cohorts reflecting real world randomized, placebo-controlled prospective clinical trials on 4 different FDA-approved drugs. *Right*: A subset of 34 (reduced model) of the 1284 genes (full model) emerged as a robust classifier of responders from non-responders regardless of cohort heterogeneity or the drug class tested. **B-C**. Bar plots show the comparison of Akaike Information Criterion (B) and Bayesian Information Criterion (C) of full model (1284 genes) and reduced model (34 genes) in all the seven training datasets. **D-G**. Lollipop plots show the fold enrichment of pathways within the 34 genes, as determined by Gene ontology (GO) biological processes (D), Gene ontology (GO) Molecular function (E), KEGG (Kyoto Encyclopedia of Genes and Genomes) pathway (https://www.genome.jp/kegg/; (F) and Reactome (https://reactome.org/; (G).

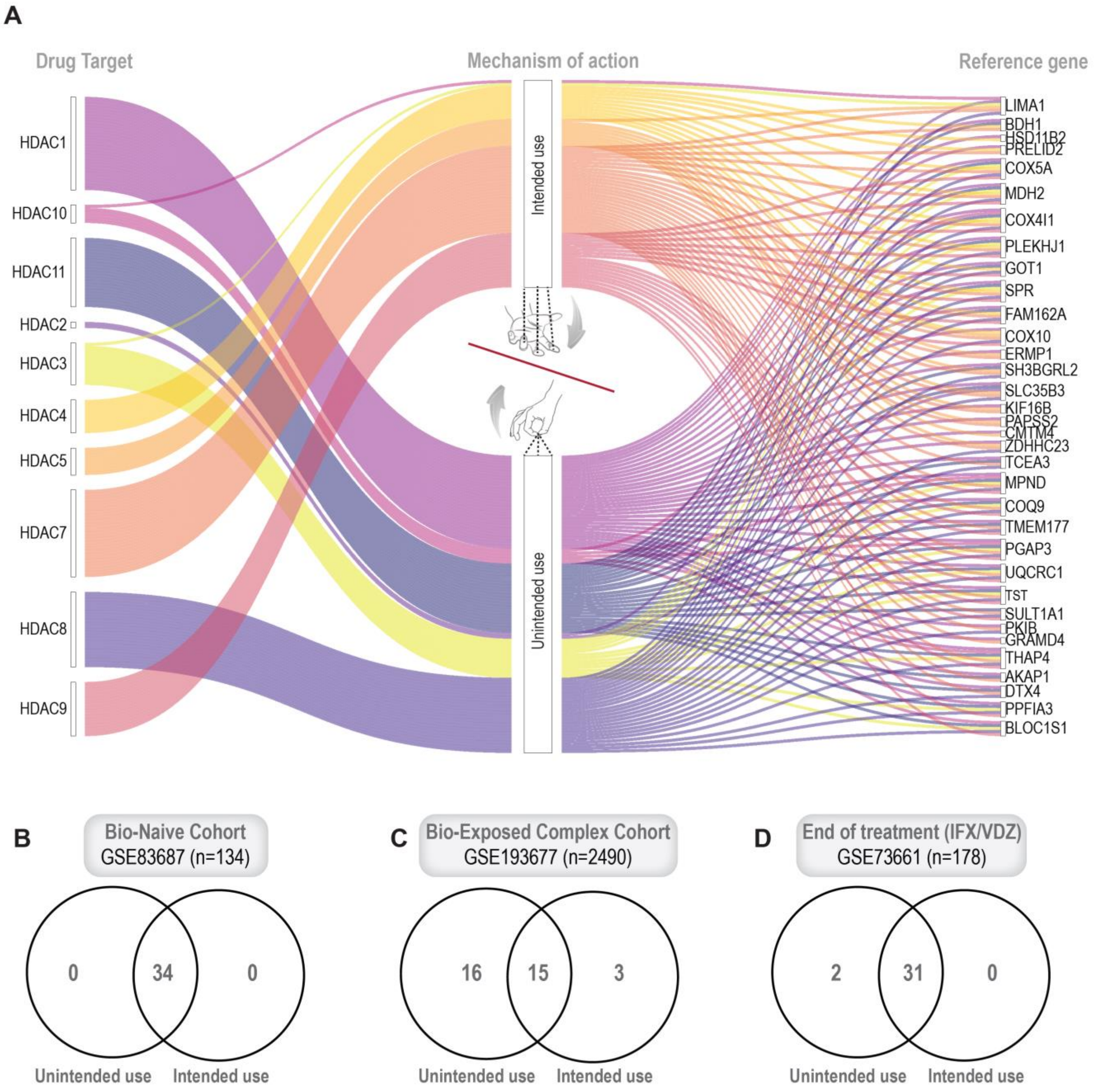


**Figure S3.** ***Likelihood of approval (LoAI) of the pan-Histone deacetylase (HDAC) inhibitor Givinostat.***
**A**. Alluvial plots for the target genes of the pan-HDAC representing the connectivity stream between target genes (*left column*) and the reference genes within the 34-gene signature of therapeutic remission (*right column*). Intended use (*top*) refers to the antagonistic action of the drug whose predictable suppression of the target gene is expected to **induce** the 34 reference genes. Unintended use (*bottom*) refers to the impact of antagonistic action of the drug whose predictable suppression of the target gene is expected to **suppress** the 34 reference genes. **B-D**. Venn diagrams of the number of 34 reference genes that are opposingly impacted in three *in silico* trials in diverse cohorts. While intended and unintended use completely negates each other in a bio-naïve cohort (A-B; the net LoAI = 0), unintended use effect is higher than intended use in cohorts that have prior exposure to biologics (C-D).

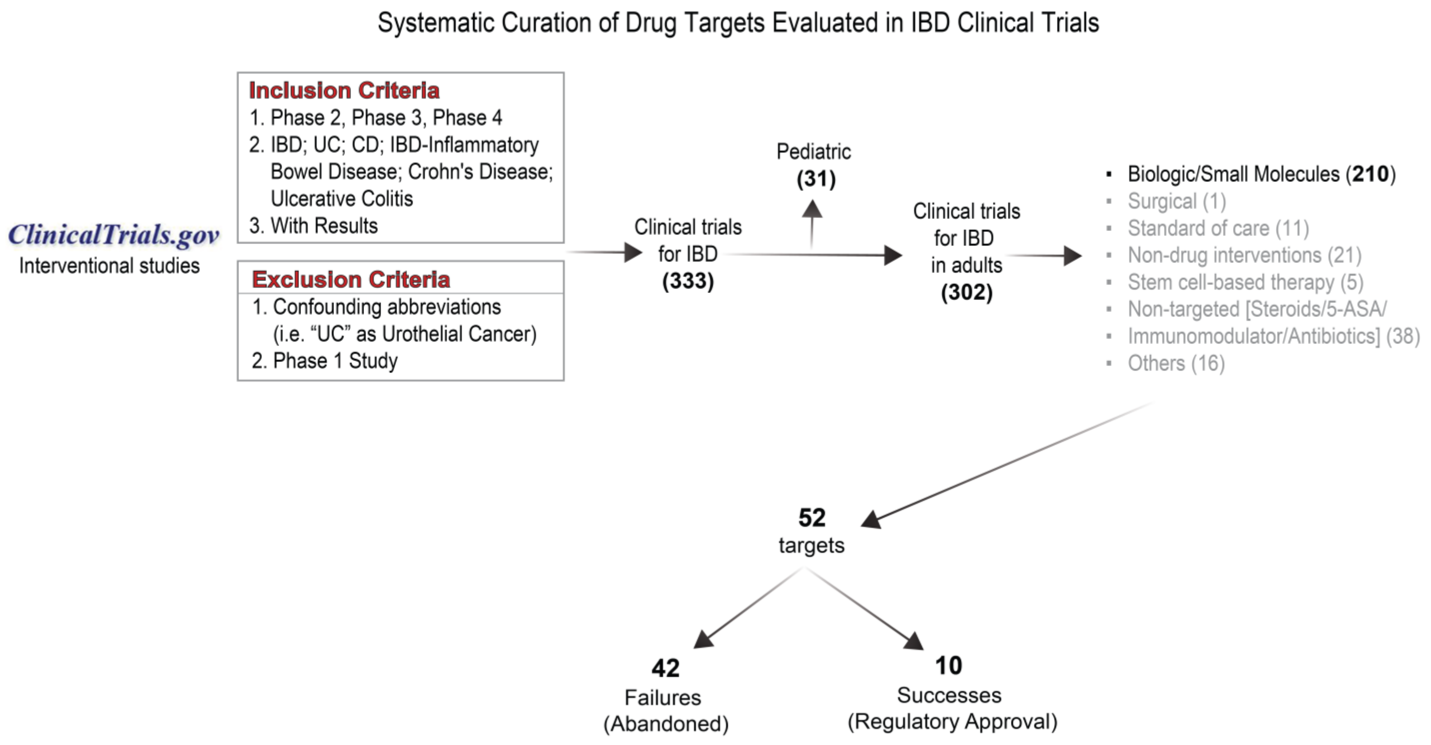


**Figure S4.** ***Systematic curation of drug targets evaluated in IBD clinical trials***
**A.** Flowchart illustrates the steps and criteria by which all the 52 drug targets were systematically chosen for evaluating the performance of F.O.R.W.A.R.D.

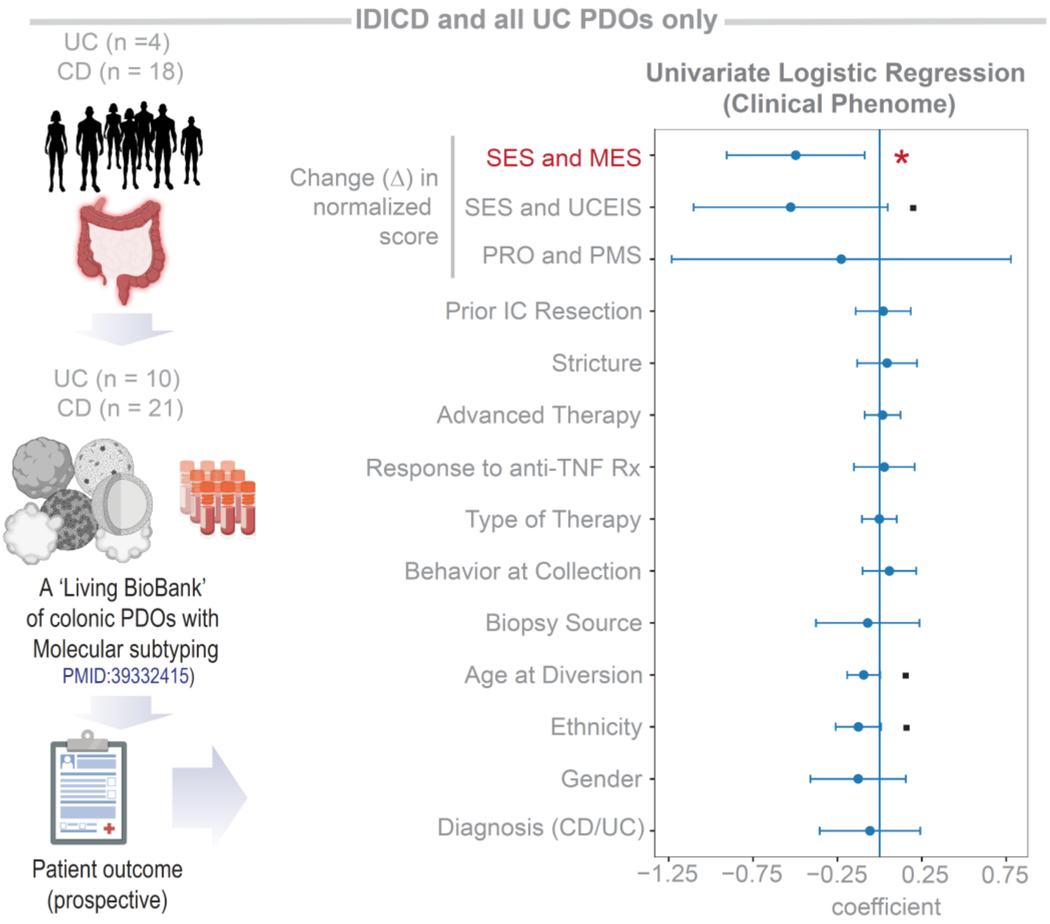


**Figure S5**. ***Univariate logistic regression analysis on IDICD and UC PDOs only.***
Univariate logistic regression model of the composite score of the 34-gene signature as a base variable, tested against each clinical covariate collected over a 5-year prospective follow-up in a previously curated IDICD/UC PDO cohort. SES, Simple endoscopic score for Crohn's disease; MES, Mayo endoscopic score; UCEIS, Ulcerative Colitis Endoscopic Index of Severity; PRO, patient-reported outcome; PMS, partial Mayo Score. The three variables on the top of the list were scaled across molecular subtypes. See also **Fig 5K** for a multivariate model on exclusively the IDICD/UC population.

| Dataset | LRT Statistic | p-value | AIC Full | AIC Reduced | BIC Full | BIC Reduced |
|---|---|---|---|---|---|---|
| GSE12251 | 0 | 1 | 5396 | 178 | 8433.45 | 252.94 |
| GSE14580 | 0 | 1 | 5398 | 180 | 8549.30 | 257.75 |
| GSE16879 | 0 | 1 | 5436 | 218 | 10147.21 | 334.24 |
| GSE73661 | -4 | 1 | 2700 | 110 | 4206.80 | 148.61 |
| GSE207022 | -2 | 1 | 5490 | 328 | 12404.38 | 495.37 |
| GSE234736 | -6 | 1 | 2514 | 156 | 4752.68 | 218.91 |
| EMTAB7604 | -12 | 1 | 2644 | 142 | 4896.55 | 201.88 |

**Supplementary Table 1:** ***Likelihood Ratio Test, Akaike Information Criterion, and Bayesian Information Criterion comparisons between full (1284-gene) and reduced (34-gene) models across training datasets.***

Likelihood ratio test (LRT) statistics comparing the full logistic regression model using all 1284 genes and the reduced model using only the 34 genes across seven independent datasets. The table includes the LRT statistic and corresponding p-value for each dataset, along with the Akaike Information Criterion (AIC) and Bayesian Information Criterion (BIC) for both the full and reduced models.

**Details of Supplemental Datasheets.** (Uploaded as separate excel files)

- **Supplemental Datasheet 1**: A catalog of all datasets and patient characteristics included in this analysis.
- **Supplemental Datasheet 2**: Connectivity between drug's target genes and the 34 reference genes of remission.
- **Supplemental Datasheet 3**: List of gene signatures used in this study.
- **Supplemental Datasheet 4**: Clinical details used in the analysis of patient derived organoids.
